# The daily life of a hummingbird: High-throughput tracking shows a spectrum of feeding and movement strategies

**DOI:** 10.1101/2025.02.25.640146

**Authors:** Jay Jinsing Falk, Alyssa J. Sargent, Jorge Medina, Alejandro Rico-Guevara

## Abstract

Most pollinators, with their small size and flight ability, are a challenge to study in the wild, yet their behavior is essential for understanding patterns of biodiversity. For example, hummingbirds play a significant role in their ecosystems—their movements from plant to plant across landscapes ultimately determines their potential as pollinators, but these behaviors are poorly understood. Two movement types are most commonly assumed in hummingbirds: territoriality and traplining, the latter strategy involving repeated and predictable visitation to dispersed feeding locations. However, direct evidence for traplining mostly comes from captive birds. In this study, we collected data from white-necked jacobin hummingbirds (*Florisuga mellivora*) that were implanted with tiny radio-frequency identification (RFID) tags, tracking their movement among a network of 20 tag-detecting feeders spread across the town of Gamboa, Panamá, for 99 days. The resulting data cover over 47,000 feeder visits from 97 freely moving birds. Overall, we found scant evidence for traplining as a consistent strategy in this species.

Instead, we identify three clusters of daily movement types, two of which are difficult to characterize as either territoriality or traplining. Our findings demonstrate that a diversity of movement strategies can be found within a single hummingbird species and even within individuals, and that many questions remain about the movement of these ecologically key vertebrates. To better understand the ecological role of hummingbirds, the description of a greater diversity of movement types beyond territoriality and traplining is likely to be necessary.

## Introduction

The movement of animals plays a key role in broad ecological phenomena such as nutrient cycling, seed dispersal, and the spread of disease (Nathan 2008). Pollinator movement dictates the extent to which many plants are able to sexually reproduce, which in turn shapes landscape biodiversity patterns (Ohashi and Thomson 2009a; Hadley et al. 2014; Wolowski et al. 2017; Betts et al. 2019). The movement of animals is shaped by a need to collect resources efficiently (Smetzer et al. 2021), and various movement “syndromes” have been hypothesized to describe these patterns (Abrahms et al. 2017, Abrahms et al. 2021). The rapid development of tracking devices in recent decades now enables high-resolution, temporally informed reconstructions of animals’ trajectories across geographic space (Kays et al. 2015; Hurme et al. 2025). However, pollinating animals like insects and hummingbirds are often small, fast-moving, and travel long distances, making them a unique challenge to track in the context of their natural habitats (Hadley and Betts 2009; Rueda-Uribe et al. 2024).

Besides their critical role in many ecosystems, hummingbirds represent an extreme in terms of vertebrate movement capability. They hold records for the fastest speed per body length (Clark 2009), can sustain rates of nearly 100 wingbeats per second (Feo and Clark 2010; Clark 2011), and perform one of the longest migrations relative to body size (Healy and Calder 2020). Despite extensive work on these flight capabilities, tracking the movement of individuals in the wild can be difficult due to their small tarsi which do not allow for color banding (Kapoor 2012). Color-coded back-tags are also difficult to use for many individuals simultaneously (Rico-Guevara and Araya-Salas 2015), and radio-telemetry tags have only recently become small enough to place on larger hummingbirds (Hadley and Betts 2009; Zenzal et al. 2014; Torres-Vanegas et al. 2019). Even these tags tend to be short-lived, or are difficult to use in many habitats (Hadley and Betts 2009; Rueda-Uribe et al. 2024).

Due to their record-high metabolic rates (Suarez 1992), hummingbirds’ movement is pronouncedly tied to their foraging strategies (Hadley et al. 2018; Hazlehurst and Karubian 2018). Previous research on these strategies has tended to focus overwhelmingly on two potential alternatives: territoriality and traplining (Sargent et al. 2021). In this context, territoriality is an interference competition strategy that involves physically excluding other nectarivores (e.g., hummingbirds, flowerpiercers, insects) from access to floral resource patches (Wolf 1970; Rico-Guevara et al. 2021; Sargent et al. 2021). Conversely, traplining is an exploitation competition strategy defined by the use of long, repeated circuits from one nectar resource to another (Janzen 1971; Gill 1988), and does not require physical exclusion of competitors at each resource (Feinsinger and Colwell 1978; Garrison and Gass 1999; Volpe et al. 2014; Rico-Guevara et al. 2021; Sargent et al. 2021). The territoriality/traplining distinction has been used to describe behavioral variation not only among species (Stiles 1975; Tiebout III 1991; Cotton 1998; Altshuler et al. 2004; Betts et al. 2015; Rodríguez-Flores and Arizmendi Arriaga 2016; Goldshtein et al. 2020; Rombaut et al. 2022), but also between sexes—males are more often described as territorialists, and females as trapliners (Sargent et al. 2021; Temeles et al. 2005).

These hypotheses and assumptions have been difficult to test without modern techniques to observe the individual movement strategies of multiple individuals over long periods of time, and, as such, many studies rely on traditional foraging classifications to simplify the analytical process (Altshuler et al. 2004; Betts et al. 2015; Rombaut et al. 2022). In this study, we used small Radio-Frequency Identification (RFID) tags (Passive Integrated Transponders or PIT tags) to track individual hummingbirds across a large array of tag-detecting feeders. We studied the movement of white-necked jacobins (*Florisuga mellivora*) in Gamboa, Panama, a small town with extensive edge habitat adjacent to the Panama Canal and surrounding lowland tropical forest of Soberanía National Park. White-necked jacobins are frequently seen engaging in aggressive territorial behaviors around feeders and flowers (Stiles et al. 2020). Females may be less territorial, but this nuance can be difficult to discern because females are polymorphic, with some looking like males (Falk et al. 2021, 2022). As with most hummingbirds, the extent to which white-necked jacobins vary in their strategies, both between individuals and over time, is unknown.

We first tested whether territorial and traplining strategies were being used and their relative extent in this population. Second, we used a descriptive method with no *a priori* assumptions of movement type and attempted to qualify movement types. Ultimately, testing assumptions of movement strategies in a large number of individuals to identify the types and forms of variants is an important step for understanding the natural history and movement ecology of a group of organisms, especially for an animal representing extraordinary feats of energy consumption, pollination ecology, and movement capability.

## Methods

Full capture, sexing, tagging, and detection methods can be found in the Supplemental Methods and in Falk et al. (2021). Between December 2017 and May 2019, we subdermally tagged white-necked jacobins with Biomark HPT8 PIT tags in and around the town of Gamboa, Panama and Soberanía National Park. We genetically sexed all birds using PCR on DNA extracted from blood, with either 2550F/2718R (Fridolfsson and Ellegren 1999) or 1237L/1278H primer pairs (Kahn et al. 1998).

Hummingbird feeders with a single feeding port equipped with a PIT tag detector were installed in fixed locations around Gamboa with at least 90 meters of separation, and no line of sight between feeders. Our detection accuracy tests in captivity showed that 100% of feeds were detected, and tags were not detected during any behavior other than feeding (Falk et al. 2021). Only one tag could be logged at a time, but antennas were oriented in such a way that having two birds near the port would not be possible. Antennas searched for tags 3 times/sec, and could detect tags within roughly 5 cm from the feeding port. Presence and absence at the antenna were logged every second, and the duration of a visit was the sum of all feeding seconds plus gaps of 7 seconds or less (See Supplemental Methods for details). Various numbers of feeders were activated between January of 2018 and May 2019. However, we narrowed our analyses toward the end of this period, between February 18, 2019 – May 27, 2019, when the number of tagged birds was at a maximum and the number of RFID feeders did not change. 107 individual birds were detected during this 99-day period.

During this period, we actively maintained and collected data from 20 feeders. We randomly chose half of the feeders to have a high sugar concentration (1:3 sucrose:water, vol:vol), and the other half to have a low sugar concentration (1:6 sucrose:water, vol:vol). Concentration assignments were switched at the middle of the sampling period (on April 8th) such that high-concentration feeders became low, and vice versa.

### Describing daily life

To describe daily movement patterns, we extracted numerical descriptors for each individual bird for each day that it was detected (henceforth referred to as a “bird-day”). We assumed that at least 3 feeder visits were necessary to include a given bird-day in the analyses, and excluded any bird-days with fewer visits. For each bird-day we extracted 6 variables: 1) total number of feeder visits, 2) mean duration of each feeder visit, 3) total number of feeder stations visited, 4) largest distance between any pair of visited feeders, 5) diffuseness, and 6) mobility. Diffuseness is a measure of spread across multiple feeder station visits (opposite of centralized movement, the high usage of a single feeder), and was calculated by dividing the number of feeds at feeders away from the bird’s “primary” feeder (the feeder from which an individual fed the most at a given day) by the number of feeds at the primary feeder. Mobility was calculated as the number of times the bird moved from one feeder to another, divided by the total number of feeds.

Providing two different sugar concentrations in the feeders (1:3 versus 1:6) was intended to introduce variation in preferred feeders, and in a previous study using the entire dataset, we found a weak but significant preference for feeders when they contained higher sugar concentrations (Falk et al. 2021). In the more limited period of this study, however, we found that in comparison to feeder location (p < 0.0001, F = 14.6), sugar concentration was not a significant factor (p = 0.76, F = 0.09) contributing to overall preference (ANOVA with both location and sugar concentration as explanatory variables for feeder visitation rate). We therefore do not include visitation to high versus low sugar feeders in our analysis of movement.

### Testing for territoriality and traplining

Movement strategies can be described by a correlation of multiple metrics. We tested for the presence of these correlations as a way to identify usage of territoriality and traplining. For territoriality, we assume that territorial birds should be detected with greater frequency within bird-days, and that bird-days with the highest number of feeder visits should be associated with lower diffuseness (high centralization to a single feeder), and with lower mobility. We evaluated these predictions using linear models.

Territorial strategies can be observed directly in white-necked jacobins simply by watching their behaviors at feeders. However, non-territorial strategies like traplining cannot be directly observed and are more difficult to infer. Classic definitions of traplining incorporate elements of *movement predictability* between specific plants or patches of resources (Feinsinger and Chaplin 1975; Ohashi and Thomson 2009b; Reynolds et al. 2013; Somanathan et al. 2019). We measured movement predictability in two ways, which we detail in the next paragraphs: 1) the similarity in movement pathways from one day to the next, and 2) within-day route repeatability (i.e., daily route determinism). If white-necked jacobins use a traplining strategy when not defending nectar resources, we expect higher measures of predictability to be associated with high diffuseness, more movement, and larger detected travel distances. We again tested these predictions with linear models.

For between-day comparisons, we compiled the pairwise transitions from one feeder to another and the number of times each pairwise transition occurred. If a bird uses a similar travel route from one day to the next, as would be expected under traplining, we should expect larger numbers of similar pairwise transitions. For each focal bird-day, we counted the number of times each potential pairwise transition between feeder stations occurred. If the bird was again present in the dataset during the next consecutive day, we calculated the absolute value of the difference in the number of each transition between the two days. We then summed all differences for all pairwise possibilities, and divided this by the total number of transitions on the focal day. Greater transition difference therefore refers to lower between-day predictability.

To calculate within-day predictability, we calculated determinism within bird-days (Ayers et al. 2015). Determinism is adapted from recurrence quantification analysis and uses recurrence plots (which quantify the number and length of recurrent sequences) to detect repeated sequences within a larger sequence of events (Ayers et al. 2015, 2018). In this case, a “recurrence” would refer to an individual hummingbird returning to a feeder that it visited previously (Ayers et al. 2015). As such, determinism is an ideal measure for detecting sequential behaviors in animals because it accounts for repeated sequences of any length, while weighting longer repeated sequences higher than shorter repeats; determinism allows for null models to test the significance of various behavioral strategies, particularly traplining (Ayers et al. 2015). We removed sequentially repeated feeds from the same feeder for each bird-day, since we regard movement between feeders as the main aspect of a route rather than repeats at the same feeder. We considered a “route/sequence” of feeds to consist of at least 3 different feeder visits, and then repeated the analysis with a minimum of 2 different feeder visits. Since the minimum number of feeder visits requires at least 1 repeat of this sequence, we removed any bird-days fewer visits than double the minimum different feeder visits (6 or 4, respectively). Reversed sequences were permitted in our calculation of determinism. We repeated the analysis to count sequences of at least 2 feeder visits for a more lenient approach. Sequence lengths of 2 indicate simple movement between 2 feeders, which is arguably not a true trapline in and of itself. However, it is possible that 2 RFID feeders are a part of a larger sequence of feeding locations that do not include our detected locations.

### Describing daily movement

To analyze bird movement patterns without *a priori* predictions, we next compiled all 6 descriptors for each bird-day, and used a principal component analysis (PCA) to identify independent axes of variation in daily movement patterns. All variables were log-transformed, scaled with variance equal to 1, and centered around the mean at 0. Results indicated that only the first two eigenvalues were greater than 1, together comprising 79.3% of daily movement variation (PC1: 57.7%, PC2: 21.6%), so proceeding analyses only used these two variables. To better understand these axes of variation, we calculated their correlations with the 6 movement variables. We plotted the first and second PC scores for each day of each individual in a scatterplot to visualize the complete “movement space” of white-necked jacobins. Using the MASS package in R (Ripley et al. 2013), we next plotted contours of a kernel density estimation onto the movement space in order to identify types of movement that were especially common or potentially distinct. To determine whether sexes differed in their movement strategies, we separated males and females to visualize their degree of overlap.

We note that when a bird was detected at a single feeder station during a given day, only the number of feeds and the feed duration can vary, while the rest of the variables all collapse to a single number: total number of stations visited is 1, decentralization and mobility are 0, and we arbitrarily assumed a movement distance of 10 meters (accounting for short flights around the feeder). This caused extensive clustering in movement space among these particular bird-days. Given normal movement stochasticity, this clustering is unlikely to be as extreme as the PCA suggests. However, the strategy and presence of these bird-days is both biologically relevant and also distinct from other movement types present in the dataset.

In addition to describing the full expanse of movement possibilities, we attempted to identify average movement types for each individual. To do so, we limited the dataset to days in which birds fed at least 3 times per day for at least 10 days. Here we chose a more stringent minimum, because too few days in the dataset may not represent an “average” movement type for any given individual. We then found the average PC1 and PC2 scores for each of these individuals. To see whether individuals fell into distinct regions of individual movement space, we then used model-based clustering methods to objectively identify the number of clusters and the cluster group for each bird using the mclust package in R (Fraley et al. 2012), which assumes the data are clustered in a variable number of Gaussian ellipsoids based on density functions, then assigns points to these clusters and calculates Bayesian Information Criterion (BIC) scores for the data matrix under 1–9 clusters (Fraley and Raftery 2007; Fraley et al. 2012). We then calculated the mean and 95% confidence interval of all 6 movement descriptors and characteristics for each group.

## Results

During the period of interest, 48,476 feeds from 107 birds were detected. After removing bird-days in which fewer than 3 feeds occurred, we retained 47,332 feeds from 97 birds, with a mean of 23.8 ± 25 SD feeder visits per day, and a total of 1994 bird-days. 30 individuals were females with 6 juveniles, and 67 were males with 6 juveniles. Among the adult females, all had the heterochromic plumage morph except for one, which was captured as a juvenile in 2018, but its plumage type could not be verified at the time of the study in 2019.

Testing for territoriality and traplining

We found a negative relationship between the number of feeds detected and both the diffuseness (estimate = -3.44, error = 0.60, *p* < 0.0001), and mobility (estimate = -14.3, error = 1.63, *p* < 0.0001). These results (Figure 2) indicate that birds that tend to use a low number of feeders, and who rarely move between feeders, are also those that feed at high rates in our dataset.

**Figure 1.**
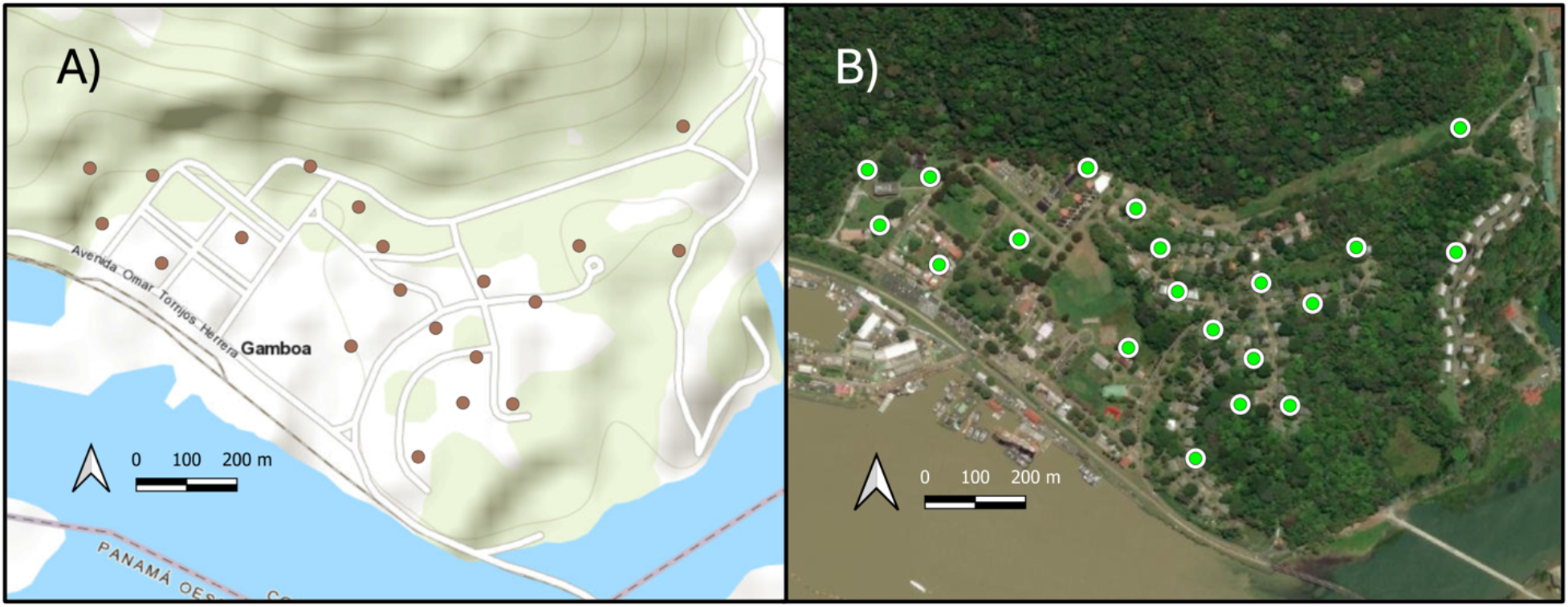
A map of hummingbird feeders with RFID detection in the town of Gamboa, Panama. A) Topographic map with feeders in brown. Gray lines indicate 20m differences in elevation. B) Satellite image with feeders in green shows that Gamboa is flanked by dense forest to the North, the Panama canal on the South and West, and the Chagres river to the East. Gamboa itself consists of edge and mixed habitat between forest and human-disturbed areas.

**Figure 2.**
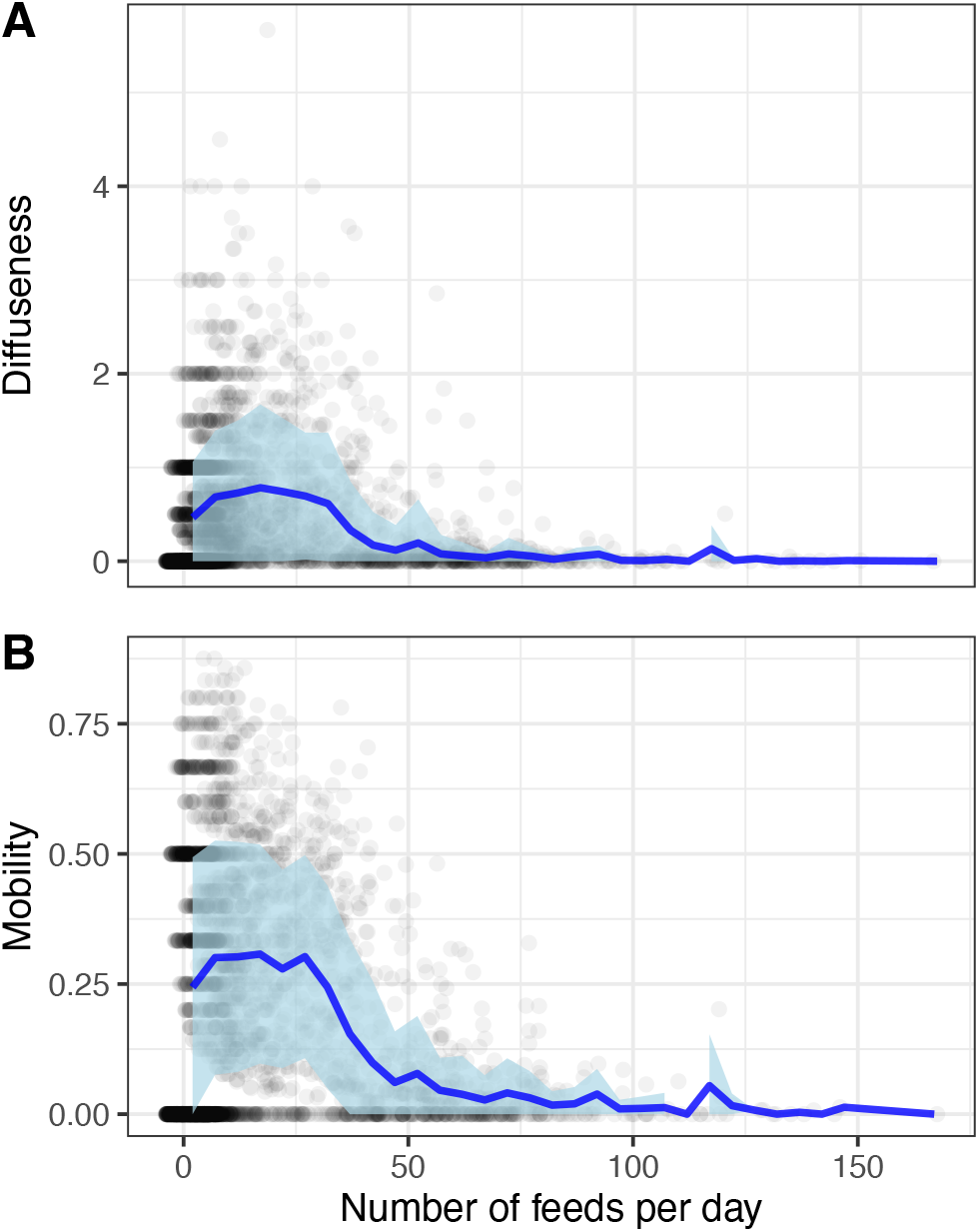
Identifying territorial strategies. Diffuseness is a measure of “spread” across feeder usage in one bird-day, while mobility is the proportion of feeds that occur at a different feeder from the previous feeder visit. Points are jittered to show data density. Dark blue lines indicate the mean across bins of 5 feeds per day, while light blue shaded regions show the standard deviation around the mean. Both (A) diffuseness and (B) mobility are on average higher and more variable at low numbers of feeds per day. At higher feeds per day, mobility and diffuseness are close to 0, consistent with a highly centralized territorial strategy where little movement is required.

We next tested for the presence of a traplining strategy. We found a positive relationship between consecutive day transition predictability and diffuseness (estimate = 0.42, error = 0.07, *p* < 0.0001) and travel distance (estimate = 131.5, error = 21.20, *p* < 0.0001). These results (Figures 3A, 3B) show that birds that are likely to be non-territorial, with high diffuseness and long travel distances, tend to be associated with lower levels of predictability across consecutive days, which is opposite to traplining expectations.

**Figure 3.**
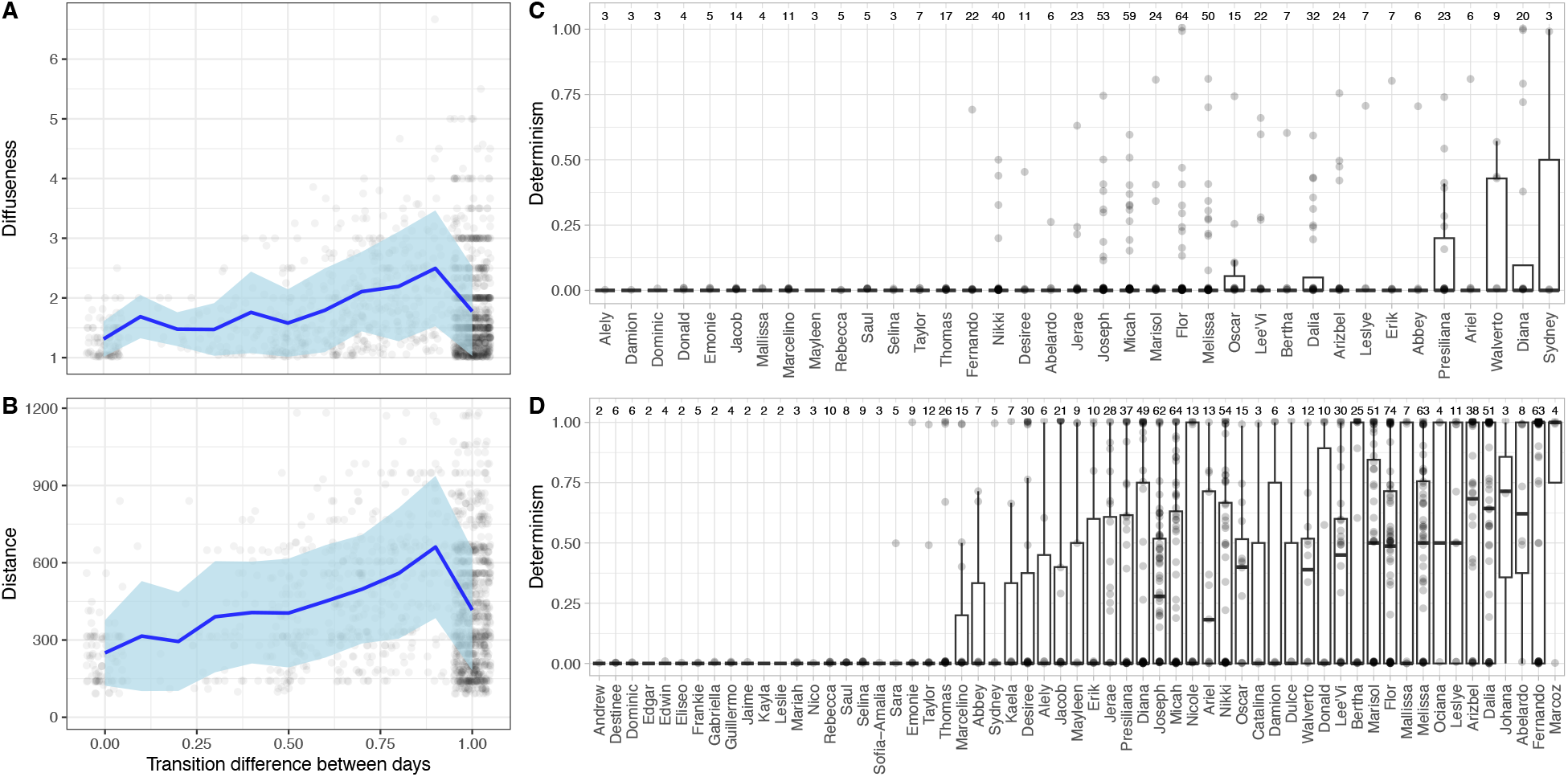
Testing for the presence of traplining. Traplining strategies should involve high diffuseness, longer travel distances, and high route predictability. (A, B) We calculated between-day route predictability as the proportion of feeder transitions that were repeated on the consecutive day by the same bird. Low transition difference between days refers to higher degrees of predictability between days. We find the opposite of what we expect for traplining strategies: long travel distances and high degrees of diffuseness were associated with lower degrees of predictability. The high degree of variation leaves space for the existence of traplining, but this does not appear to be the norm. (C,D) We calculated determinism as a measure of predictability of movement within days. Determinism is the number of route sequences that are repeated within the same day in proportion to the total number of route sequences traveled. Points, which are vertically jittered to avoid overlap, indicate determinism levels for each day in which there was sufficient data. Dark horizontal lines show the median level of determinism for each bird, whiskers show the 75% percentile, and whiskers extend to 1.5 times the interquartile range. Numbers on top indicate the number of days with sufficient data for each individual. (C) At least 3 feeders are required for a sequence to be counted. All individuals displayed low levels of determinism, with median determinism at 0 for all individuals. (D) Sequences of just 2 feeders are allowed for a more liberal approach to identify determinism. With this approach we see a greater range of determinism within and between individuals. Some individuals show higher rates of determinism than others.

We calculated determinism within days as a second measure of predictability. We found low levels of determinism in the population (Figure 3C, 3D). When the minimum feeder route length was 3, no individual with sufficient data had median levels of determinism above 0 (Figure 3C), indicating very low levels of route predictability. Higher levels of determinism were occasionally exhibited by some individuals, but not on a regular basis. We repeated the analysis with a minimum feeder length of just 2 (Figure 3D), and this time found that birds varied in their degree of determinism, with some individuals showing high levels of determinism on several days. Using determinism levels from this analysis, we found that higher levels of determinism were associated with lower space use (estimate = -34.9, error = 17.16, *p* = 0.042) and lower diffuseness (estimate = -0.18, error = 0.06, *p* = 0.002), again opposing the expectations of a traplining strategy.

### Describing Daily Movement

Principal component analysis of all bird-days in the dataset showed that PC1 is correlated with a larger number of visited feeders (91.4%), longer distances detected between visited feeders (86.9%), greater diffuseness (90.8%), and higher mobility (93.6%). PC2 is correlated with greater number correlated with greater number of feeds (80.1%) but lower mean duration feeds (−69.5%) (Table S1 for complete PC loadings and correlations). Kernel density estimation showed regions of increased density (Figure 4). Two clusters fell in a region of movement space with low PC1 (Figure 4A, left of dotted line). All points in these two regions consisted of daily behavior in which birds only visited a single feeder. As discussed previously, the clear divide denoted by a dotted line is an artifact of our methods. However, the two regions divided by the dashed line cannot be explained by a method artifact, demonstrating movement types with either high or low PC2.

**Figure 4.**
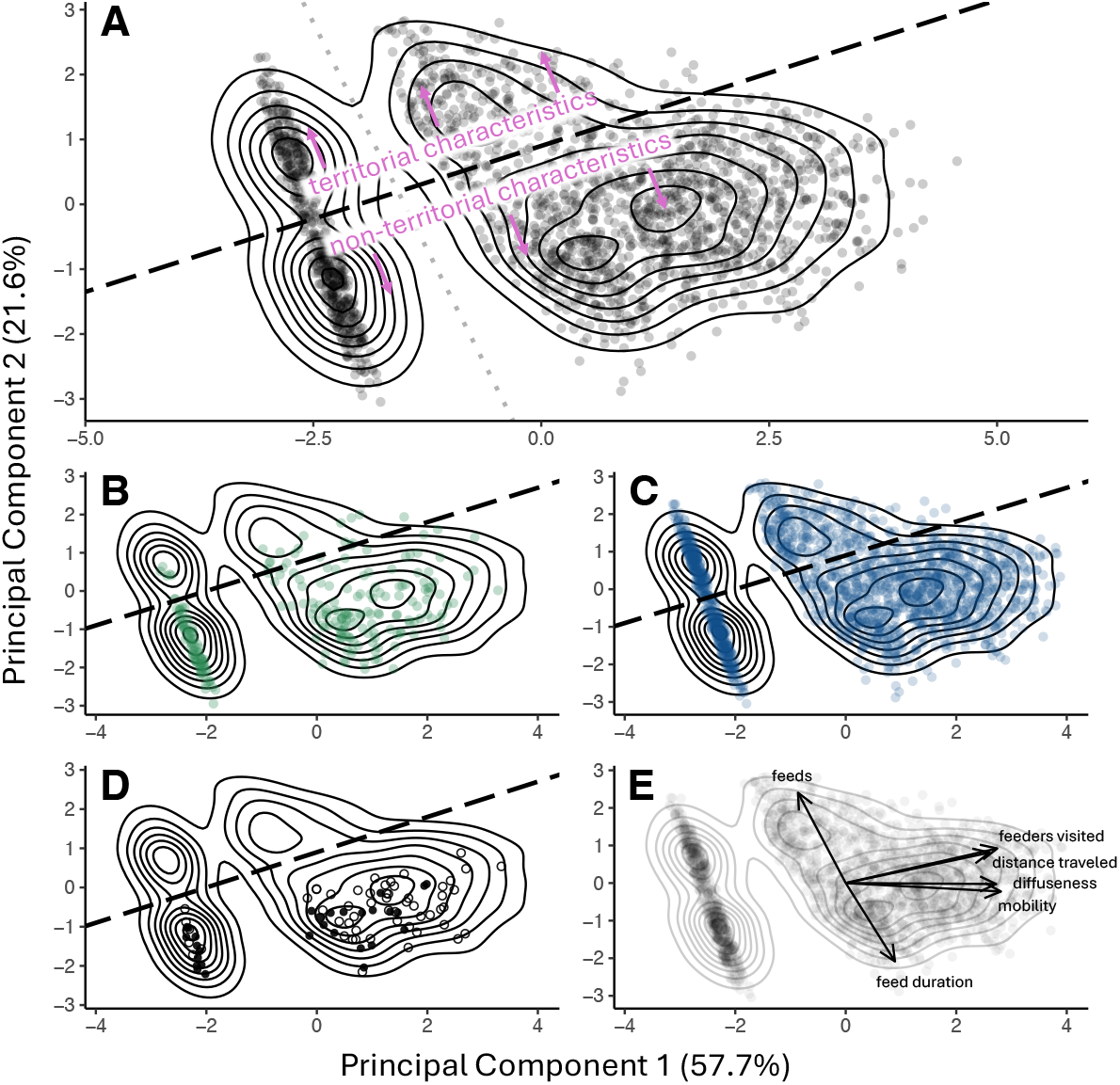
Visualizing the movement space of white-necked jacobins. A) The first two principal components of a PCA which included all 6 movement descriptors, covering 1983 bird-days. The resulting graph is a visualization of movement variation in this species. The dotted line indicates an artificial separation of movement types created because several movement parameters are invariable when a bird only visits a single feeder. Other density peaks cannot be explained by artifacts, and the dashed line shows a separation between putative territorial and non-territorial movement types. B) Only female bird-days shown. C) Only male bird-days shown. D) Only juvenile bird-days shown; females with filled circles, males with hollow circles. E) Biplot showing the loadings of each variable on the movement space with all bird-days.

### Describing regions of the PCA space

Regions of movement space were not completely separate but provide relevant topography that may represent varying aspects of a spectrum of movement types. We calculated the average number of feeds per day and the average distance traveled above and below this line, and found that individuals above the dotted line fed an average of 55.3 times per day and traveled 207.2 m between feeders, while those below the line fed an average of 12.1 times per day (t-test: t = 37.4, degrees of freedom = 606.5, p < 0.0001) and traveled 349.1 m (t-test: t = -11.3, degrees of freedom = 1105.2, p < 0.0001). This negative relationship between these distances and feeding rates should be expected if movement varies along a spectrum from territorial to non-territorial characteristics. The topography of movement space in white-necked jacobins therefore shows properties that are consistent with such a spectrum. Panels B and C of Figure 4 show the same graph but with only female and male bird-days, respectively. Both sexes exhibit more movement below the line (non-territorial characteristics), but males exhibit bird-days above the line (territorial characteristics) in greater proportion (29.2% of male bird-days and 10.0% of female bird-days are above the line, *p* < 0.0001, χ^2^= 40.7). 31.8% of adult females and 41.0% of males were above the line for at least one day. Among these individuals, males tended to be more consistently above the line than females (40.0% versus 19.2% respectively). Overall, neither sex exhibited putative territoriality with consistency, but males tended to do so more often than females and juveniles (juveniles never exhibited putative territoriality).

### Individual movement space

We next averaged the PC1 and PC2 scores for each individual with over 10 days present in the dataset, resulting in a total of 47 individuals (mean days 37.4 ± 29.6 SD, total range 11–99 days, Figure 5). 39 were males with 3 juveniles, and 9 were females with 1 juvenile. The highest BIC score from model-based clustering came from dividing the movement space into 3 ellipsoid clusters (−235.9, 0.82 higher than the next best model with 2 clusters). Class 1 contained 31 individuals while the other two were smaller and contained 9 and 8 individuals (classes 2 and 3, respectively). Class 1, the largest group, is characterized by a lower number of detected feeds and higher feed duration (Figure 5). Class 2 was distinguished by a large number of feeders visited, high diffuseness, and higher space use (Figure 5). Class 3 was characterized by the highest number of overall detected feeds per day, lower feed duration, as well as lower distance traveled, number of feeders visited, mobility, and diffuseness (Figure 5). Clusters did not significantly differ in the proportion of females and males (*p* = 0.17, χ^2^= 3.30), but sample size for females was small (n = 9). We note the number of clusters was sensitive to the minimal number of days allowed for an individual to be present in this analysis (Supplemental Table S1). In some of these cases, Classes 1 and 2 would be clustered together, but Class 3 was nearly always distinct, causing there to be 2 rather than 3 clusters.

**Figure 5.**
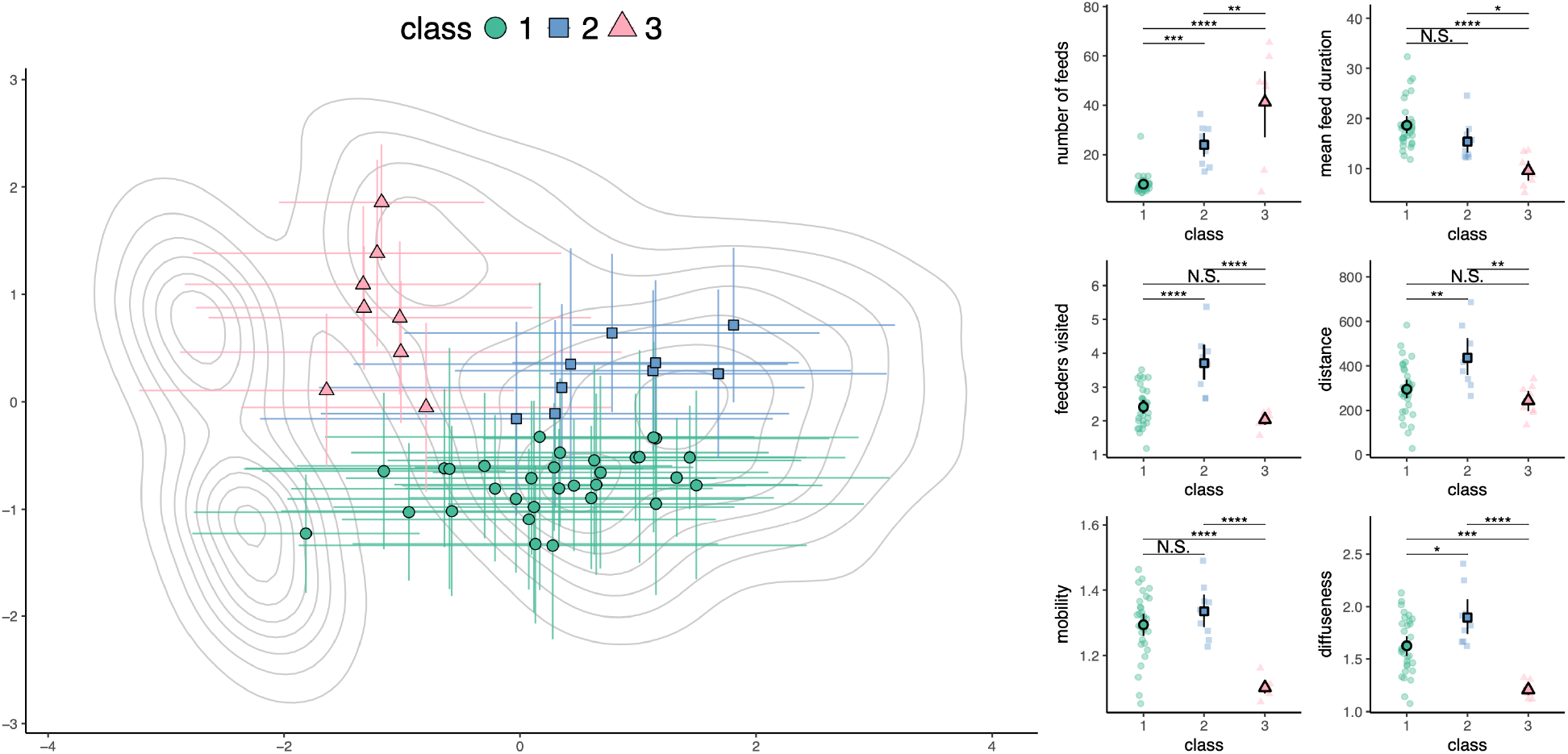
Individual strategies of white-necked jacobins. Movement metrics were averaged for 47 individuals with at least 10 days in the dataset. Cluster analysis separated individuals into 3 different classes. On the left panel, mean movement for each individual is shown overlaid on top of the density contours of all bird-days (i.e., Figure 4A). Lines extended from points show standard deviations around the mean for both axes. Coloration and shape designate each of the 3 classes. On the right we show how each of the 3 classes differ in the 6 movement types. Means are shown with bootstrapped 95% confidence intervals. p-values for differences in means are indicated by N.S.: p > 0.05, *: 0.05 > p > 0.01, **: 0.01 > p > 0.001, ***: 0.001 > p > 0.0001, ****p < 0.0001.

## Discussion

Pollination ecology relies on accurate theory on the movement of animal pollinators, yet pollinators are often small and difficult to observe in their natural habitats. Movement of these pollinating animals, hummingbirds especially, are often described as either territorial or traplining (Feinsinger et al. 1979; Lemke 1984; Gill 1988; Tiebout III 1991; Cotton 1998; Volpe et al. 2014). Corroborating our direct observations of white-necked jacobins, we identified bird-day descriptors which align with a territorial strategy. A high number of short visits to a small number of centralized feeders, sometimes exceeding 100 visits per day (Figures 1 and 5), is most likely to be associated with the territorial behavior seen in the wild.

Most daily behavior patterns in the data are easier to describe as non-territorial, yet evidence that non-territorial strategies involve traplining as a consistent strategy was scant. Definitions of traplining vary, but all entail a degree of route predictability over many feeding areas, often implicitly or explicitly suggesting coverage of larger total areas (Ackerman et al. 1982; Williams and Thomson 1998; Gass and Garrison 1999; Lihoreau et al. 2013; Somanathan et al. 2019; Torres-Vanegas et al. 2019). Traplining strategies are also diffuse, since individuals are expected to move from location to location along a route rather than defending a single location. We employed two measures of route predictability between feeders, one covering the predictability across consecutive days, and the other within days. Across consecutive days, we found that increased predictability in feeder transitions was associated with lower distance coverage, and lower diffuseness (Figures 3A, 3B), in contrast to the predictions of a traplining strategy. Route determinism more than 0 was rare when routes were restricted to 3-feeder minima (Figure 3C), and only appeared in 15% of the 620 bird-days with sufficient data, and only 4% showed determinism levels above 0.5. No individuals showed median levels of determinism greater than 0. In other words, repeated sequences of 3 feeders or more are difficult to encounter on any given day, and do not appear to be a consistent strategy. We further relaxed restrictions by reducing the minimum route length to 2 feeders (Figure 3D). While 2 feeders may not be considered a “route,” due to the nature of an RFID grid and our inability to detect feeds away from our RFID feeders, we tested this more lenient definition as well. By doing so, we found many more instances of determinism, indicating that 2-feeder repeats are much more common than 3-feeder repeats. However, like we found previously, highly deterministic bird-days also involved lower diffusion and distance coverage, contrary to definitions of traplining. Rather than showing traplining, higher degrees of determinism in this case likely occur from simple back and forth movement between nearby feeders (e.g. Figure 6E). In fact, days with the highest determinism level of 1 were more likely to fall within regions of movement space (Figure 4A) associated with territorial characteristics than non-territorial characteristics (χ^2^ = 65.7, *p* < 0.0001), indicating that highly predictable movement strategies tend to have more similarity with territoriality than traplining.

**Figure 6.**
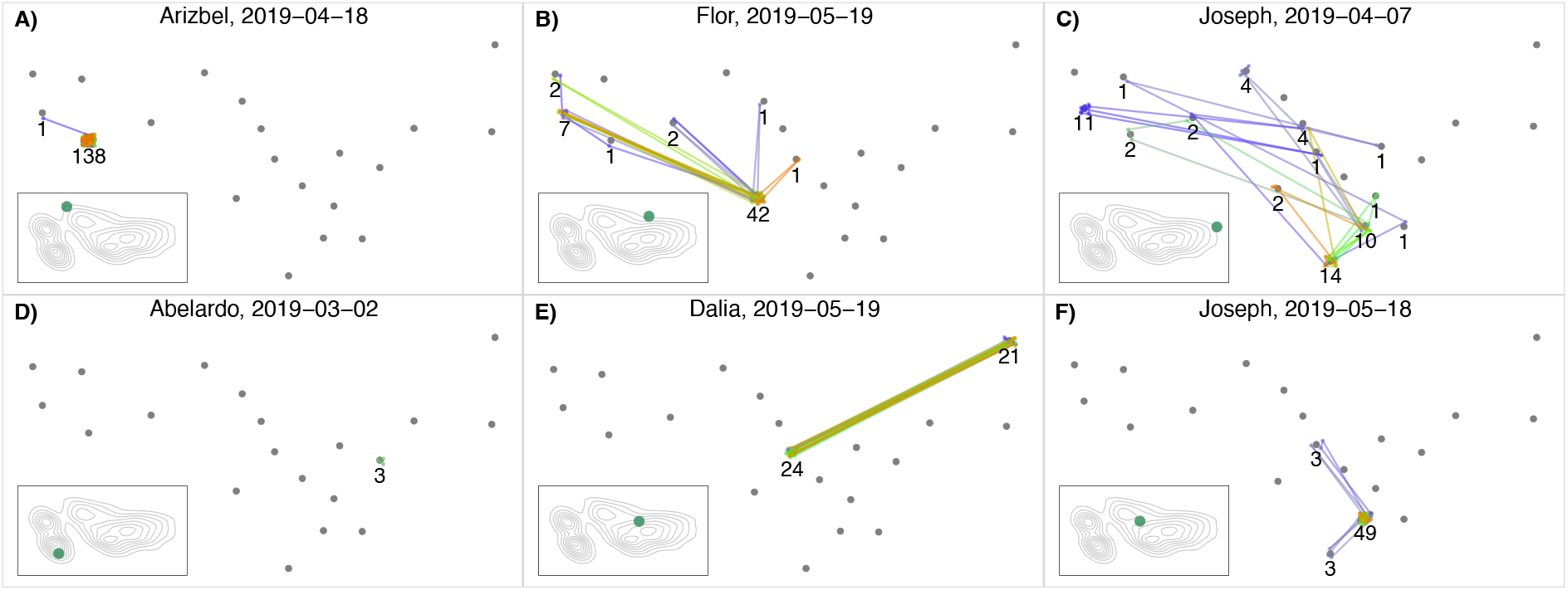
Example daily movements among RFID feeders of the white-necked jacobin. A single day’s movement is shown across 20 feeders. Bird name, which is randomly assigned without regard to sex, as well as date are shown on the top of each panel. Grey points indicate available antenna-equipped RFID feeders. Points show feeds; lines show movement between feeders. Color spectrum indicates time of day, with purple representing mornings, green midday, and red late afternoon. Number of total daily visits to each feeder is indicated. Insets show the placement of each day within movement space (i.e., Figures 4 & 5). A) High number of feeds with centralization likely indicates territoriality at a single feeder. B) Highly centralized movement involving possible exploration to nearby feeders throughout the day. C) Diffuse movement throughout the day over a wide range of feeders, and with little predictability of movement may indicate diffuse exploration or “marauding” behavior. D) Few feeds at low number of feeders indicate that a bird must be using undetected feeding locations. However, these low numbers of visits to the feeders are often longer in duration, demonstrating a possible biological relevance rather than simply missing data. E) Repeated movements back and forth between two feeders may be part of a simple trapline. F) Predictable movement between multiple feeders in the morning appears to be a small trapline, but is combined with highly centralized, possibly territorial, behavior throughout the rest of the day. Individuals can therefore shift rapidly between strategies.

Since being coined by Janzen (Janzen 1971), traplining has been discussed extensively as a movement strategy for pollinators (Thomson 1996; Saleh and Chittka 2007; Ayers et al. 2018), and the presence of traplining is believed to have impacts on plant genetic variation and ecology (Torres-Vanegas et al. 2019; Leimberger et al. 2022). Traplining is also thought to increase pollinator foraging efficiency by maximizing energy rewards and reducing search effort (Ohashi et al. 2007; Buatois and Lihoreau 2016; Klein et al. 2017; Tello-Ramos et al. 2019).

Despite this, direct evidence of traplining in wild hummingbirds is surprisingly scant. Instead, traplining is largely inferred based on repeated visitation to a single location without resource defense (Stiles 1975; Temeles et al. 2006), or long-distance daily movement exhibited by individuals (Stiles and Wolf 1979). The best evidence of feeding predictability in hummingbirds comes from observations either in captivity, or with food resources no longer than a few meters apart (Tello-Ramos et al. 2015, 2019, 2022), but there is less information from long-distance routes (see Sargent et al. 2021 for a more detailed discussion). Our data indicate that predictable routes, when present, do not necessarily coincide with long-distance foraging.

Predictable visitation rates can arise through many movement strategies which are not limited to traplining. Indeed, birds that are mostly territorial may visit alternative undefended resources at predictable intervals, giving the appearance of a trapliner if observed only from the undefended resource. In other words, while territoriality and traplining are often viewed as dichotomous and opposite to each other, we strongly caution against the synonymizing of traplining and non-territoriality without further evidence from wild animals. By avoiding this conflation, movement and pollination ecologists can move beyond idealized behaviors to identify more accurate concepts and descriptions for pollinator movement theory (e.g. Goldshtein et al. 2020).

Rather than testing *a priori* assumptions about movement syndromes, our dataset allows us to gain a fuller understanding of the spectrum of white-necked jacobin movement with quantitative methods. Our first analysis looked at the full spectrum of nearly 2000 daily hummingbird movements, mapping the “movement space.” We found two general regions of movement strategy (Figure 4). The smaller of the two spaces is generally in line with territorial strategies, with frequent, shorter feeds across smaller distances, fewer feeders, and lower diffuseness and movement. Both sexes more often exhibited non-territorial movement space (Figures 4B,C), but males tended to use territorial movements more often than females. To identify consistent strategies used by individuals, we averaged PC scores and identified 3 clusters of strategies. Class 3 is perhaps the most distinguishable based on low feed durations and higher number of daily feeds. These birds seemed to have consistently used a more territorial strategy, with lower feeder durations likely due to their continuous access to food.

Class 1 was distinguished by low detection rates at a small number of feeders but with visits of longer duration, while Class 2 individuals were detected at a larger number of feeders, and were associated with higher degrees of mobility and distance coverage.

Classes 1 and 2 would be difficult to describe as territorial, and do not consistently show signs of traplining. How else could these movement syndromes be described? Feinsinger and Colwell (1978) proposed that hummingbird movement could be described by 5 different movement strategies: high-reward trapliners, low-reward trapliners, territorialists, territory “parasites” (either large marauders or small filchers), and generalists that use multiple strategies. White-necked jacobins were listed as potential marauders, large and agile territory parasites that can move between, and swiftly feed from, rewarding flower patches even when defended by territorialists. While we did occasionally witness behavior that might align with this description, there has been little to no development of this concept, and it is difficult to know how we should expect it to manifest in movement and feeding data. Speculatively, Class 2 individuals (Figures 5, 6C), with their presence at many feeders across the landscape and high degrees of mobility, could be territorial parasites. More importantly, our data show that, as a species, white-necked jacobins can vary considerably in their behaviors, both between and within individual daily behavior. While this points to a somewhat generalist type of behavior, it is difficult for us to know whether this species is more generalist than others without similar datasets in other species.

One particularly intriguing movement type was what appeared to be exploratory movement. For example, in Figure 6B, we see a bird that uses a single primary feeder throughout the day, but repeatedly seems to venture away, visiting other nearby feeders before returning to the primary feeder. In Figure 6C, we see less centralization and less predictable movement patterns. In many cases (e.g., Supplemental Figure S2) we saw these more chaotic movement patterns—involving several feeder visits over longer distances—eventually simplify to many feeds at a single feeder (possible territoriality, e.g., Figure 6A), but continuing with centralized and nearby visits (e.g., Figure 6B). Exploratory behavior should be expected to some degree in all pollinating animals, since nectar resources are typically ephemeral and can change rapidly through the seasons (Zimmerman and Cook 1985; Lihoreau et al. 2010; Sargent et al. 2021). Obtaining knowledge of alternate resources may be critical to survival even if it appears to be less efficient in the short term. Exploratory behavior may also be important from a pollinating plant’s perspective (Torres-Vanegas et al. 2023), since long-distance pollination may occur by territorial and non-territorial birds alike (Schulke and Waser 2001; Jacobi and Antonini 2008).

As a tool for describing movement syndromes, RFID differs from many traditional methods such as GPS and radio-tracking (Abrahms et al. 2021). Chief among these differences is that RFID antennas are highly localizable and accurate at the expense of detection range.

This allows us to identify a precise behavior (feeding) during detection, and placement of multiple detection antennae can broaden our ability to detect movement. However, it is important to remember that we have no knowledge of a bird’s behavior when it is away from our 20 RFID feeders, which limits our ability to define movement types with complete confidence.

For example, in this study, we suspect certain movement types are associated with territorial behavior, based on observations of behavior in the wild, and theoretical patterns such as a tradeoff between the number of feeds and diffuseness of movement. However, with RFID alone we are unable to test whether these birds were directly excluding other birds’ access to feeders via aggressive behavior.

Though we set a minimum number of feeds per day to deduce movement syndromes, birds feeding at or near our minimum threshold must be feeding primarily at undetected locations in order to survive. The fewer detected feeds per day, the lower our ability to detect a true movement syndrome. Therefore, we are most confident about individuals that either visited feeders many times, or that visited many feeders (i.e., Figure 5, classes 2 and 3). Few detections during a particular day may indicate randomly low levels of detection, but some individuals may prefer non-feeder food sources, reflecting a biological relevance to low detection rates. We suspect the latter for two reasons. First, many individuals had consistently low detection rates throughout the experiment, reflecting a possible avoidance of RFID feeders, preference against them, or inability to access them. More importantly, there is an overall inverse relationship between the number of feeder visits per day, and the mean feed duration (estimate = -0.12, error = 0.01, *p* < 0.0001), and individuals with low detection rates tended to spend longer durations at feeders (Figure 5). Feed duration is perhaps the most reliable of our 6 movement metrics because we have complete knowledge of visitation length during each feed. Since feed duration has a non-random association with detection rate, it suggests that low detection rate is a relevant metric to a broader feeding syndrome rather than simply a lack of data from other strategy types. Perhaps birds that feed less frequently can do so precisely because of these longer feeding times. The difference between a long and short feed can be significant—while the lowest 1% of feed durations were only 1 second, the highest 1% averaged 75 seconds.

Finally, as with any characterization of behavior, we note that the range of movements mapped here are particular to the study at hand, and may differ in other contexts, including various habitats, landscapes, environmental conditions, and the relationship of this species to other species in this environment (Feinsinger and Colwell 1978). In Gamboa, during the dry season, white-necked jacobins are extremely common and able to chase away most other species at feeders (e.g., *Amazilia tzacatl, Phaethornis longirostris*, and *Amazilia edward*), but this is not the case in other locations (Fernandez-Duque et al. 2024). Our detection method may have also affected movement behaviors since hummingbird feeders are an especially high-reward artificial food source. Such a food resource may decrease the probability of using non-territorial behaviors such as traplining because the payoff for resource defense would be very high. Nevertheless, we did found extensive evidence for non-territorial behaviors. Our hope is not to deny the existence of traplining, but simply to emphasize that traplining does equate to non-territoriality, and that non-territoriality likely takes many forms.

Detecting movement with RFID has pros and cons, but the method remains one of the best ways to detect long-term animal movements from a large number of individuals, especially when the animals are small. Our dataset represents one of the most comprehensive studies of hummingbird movement, yet we found little evidence for traplining. This movement type may exist in other species, but the scant evidence from wild hummingbirds should give pause to the assumption that when hummingbirds are not behaving territorially, they are regularly engaging in traplining. Non-territoriality may in fact take numerous forms, but adherence to the trapline ideal limits our ability to identify real pollinator movement strategies, and may perpetuate inaccurate narratives on the flow of pollen across landscapes. Ultimately, full descriptions of wild pollinator behaviors may require a combination of RFID and other techniques such as active radio-tracking, the tags for which are quickly becoming small enough to safely study the smallest birds. Moving beyond theoretical assumptions and into a more quantitative understanding of pollinator behavior will not only settle long-standing dilemmas but also cement new bases for future questions that advance the field in fresh directions.

PERMISSIONS: Procedures were approved by Institutional Animal Care and Use Committees (IACUC) at both Cornell University (2009-0105) and the Smithsonian Tropical Research Institute (2015-0618-2018-A1, 2016-0120-2019, 2017-0116-2020). Blood sample collections were permitted by the Panamanian Ministry of the Environment under SE/A-84-14, SE/A-11-16, SE/A-15-17, SE/A-100-17, and SE/APHBO-7-18. Blood samples were exported from Panama to the U.S.A under C.I.T.E.S. permits SEX/A-101-15, SEX/A-111-16, SEX/A-73-17, SEX/A-47-2018, SEX/A-36-19, and U.S.D.A. Import Permit 52686.

## ACKNOWLEDGEMENTS

This work was funded in part by a Smithsonian Predoctoral Fellowship. We thank Pedro Caballero, Mia Larrieu, and Jorge Garzón for valuable field assistance. Gregg Cohen and Rachel Page for technical guidance, and use of lab materials and space. The Ministerio de Ambiente Panamá for the maintenance of conserved parklands, and for permission to record our observations.

## DECLARATION OF INTEREST

We claim no conflicts of interest.

## DATA AVAILABILITY

Data and code will be published following the peer-review process.

## Notes

### Competing Interest Statement

The authors have declared no competing interest.

