## Supplemental Materials for "The daily life of a hummingbird: High-throughput tracking shows a spectrum of feeding and movement strategies"

Supplemental Table S1: We averaged principal component scores across all days of all individuals to see if movement strategies could be identified by "clustering" of mean daily movements. Once mean PC1 and PC2 scores were calculated, we used a model-based clustering method, the mclust package in R (Fraley et al. 2012), to identify the best number of clusters, and the cluster identity of each individual. To do so, a minimum number of days for each individual needed to be set so that strategies were not based on spurious movement strategies. However, higher cutoff minimums removed increasing numbers of individuals, and we found that the best number of clusters was somewhat sensitive to the minimum cutoff because higher. The number of individuals for minimum cutoffs from 1-15 days are presented with the 3 highest BIC scores, covariance structure, and the number of clusters identified. This clustering method found either 2 or 3 clusters as the best scoring model, and individuals were similarly classified across cutoff minimums when the same number of clusters were identified. In the manuscript we present the 3 classes with a minimum of 10 days per individual to balance sufficient sample size for each individual while also maintaining sufficient individuals to classify. However, this table shows that a 2-cluster categorization is also possible.

| Min days per individual | Number of individuals | Highest model BIC value, covariance structure, # of classes | 2nd highest model BIC value, covariance structure, # of classes | 3rd highest model BIC value, covariance structure, # of classes |
|---|---|---|---|---|
| 1 | 97 | -530.0, EEE, 2 classes | -532.8, VEE, 2 classes | -534.0, EVE, 2 classes |
| 2 | 87 | -437.7, EEE, 2 classes | -438.3, EEV, 3 classes | -438.9, EEV, 2 classes |
| 3 | 81 | -394.6, EEV, 2 classes | -397.3, EVV, 2 classes | -397.5, EVE, 2 classes |
| 4 | 73 | -356.3, VEV, 2 classes | -357.5, EEV, 2 classes | -360.2, EVV, 2 classes |
| 5 | 65 | -310.2, VEV, 2 classes | -310.7, EEV, 2 classes | -312.9, EVE, 2 classes |
| 6 | 62 | -297.4, VEV, 2 classes | -297.7, EEV, 2 classes | -299.9, EVV, 2 classes |
| 7 | 60 | -291.9, VEV, 2 classes | -292.5, EEV, 2 classes | -294.3, EVV, 2 classes |
| 8 | 56 | -270.2, VEV, 2 classes | -270.5, EEV, 2 classes | -271.5, EVV, 2 classes |
| 9 | 52 | -240.2, EEV, 3 classes | -251.3, EVE, 3 classes | -251.7, VEV, 2 classes |
| 10 | 48 | -235.9, EEV, 3 classes | -236.7, VEV, 2 classes | -236.8, EEV, 2 classes |
| 11 | 47 | -228.7, EEV, 3 classes | -231.0, EVE, 3 classes | -231.5, VEV, 2 classes |
| 12 | 44 | -211.9, EVI, 3 classes | -212.0, EEV, 3 classes | -213.5, EEV, 2 classes |
| 13 | 41 | -203.7, EEV, 3 classes | -204.2, EVI, 3 classes | -205.3, VVI, 2 classes |
| 14 | 38 | -193.1, VVI, 2 classes | -193.6, EVI, 3 classes | -193.6, EEV, 3 classes |
| 15 | 36 | -178.8, VVI, 2 classes | -178.9, VEV, 2 classes | -179.3, VVV, 2 classes |

Supplemental Table S2: The proportion of variance and standard deviation by each PC from the principal component analysis of all bird-days.

| | PC1 | PC2 | PC3 | PC4 | PC5 | PC6 |
|---|---|---|---|---|---|---|
| Standard Deviation | 1.86 | 1.14 | 0.85 | 0.58 | 0.35 | 0.27 |
| Proportion of Variance | 0.58 | 0.22 | 0.12 | 0.06 | 0.02 | 0.01 |
| Cumulative Proportion | 0.58 | 0.79 | 0.91 | 0.97 | 0.99 | 1.0 |

Supplemental Table S3: Shows the loadings for each PC from the principal component analysis of all bird-days.

| | PC1 | PC2 | PC3 | PC4 | PC5 | PC6 |
|---|---|---|---|---|---|---|
| Distance | 0.87 | 0.27 | 0.04 | 0.40 | 0.02 | 0.14 |
| Number of Feeds | -0.29 | 0.80 | 0.49 | -0.16 | -0.07 | 0.04 |
| Mean Duration | 0.30 | -0.70 | 0.65 | 0.02 | -0.01 | 0.01 |
| Number of Feeders | 0.91 | 0.31 | 0.12 | 0.07 | 0.08 | -0.21 |
| Mobility | 0.94 | -0.07 | -0.15 | -0.13 | -0.28 | -0.00 |
| Diffuseness | 0.91 | -0.01 | -0.07 | -0.37 | 0.17 | 0.09 |

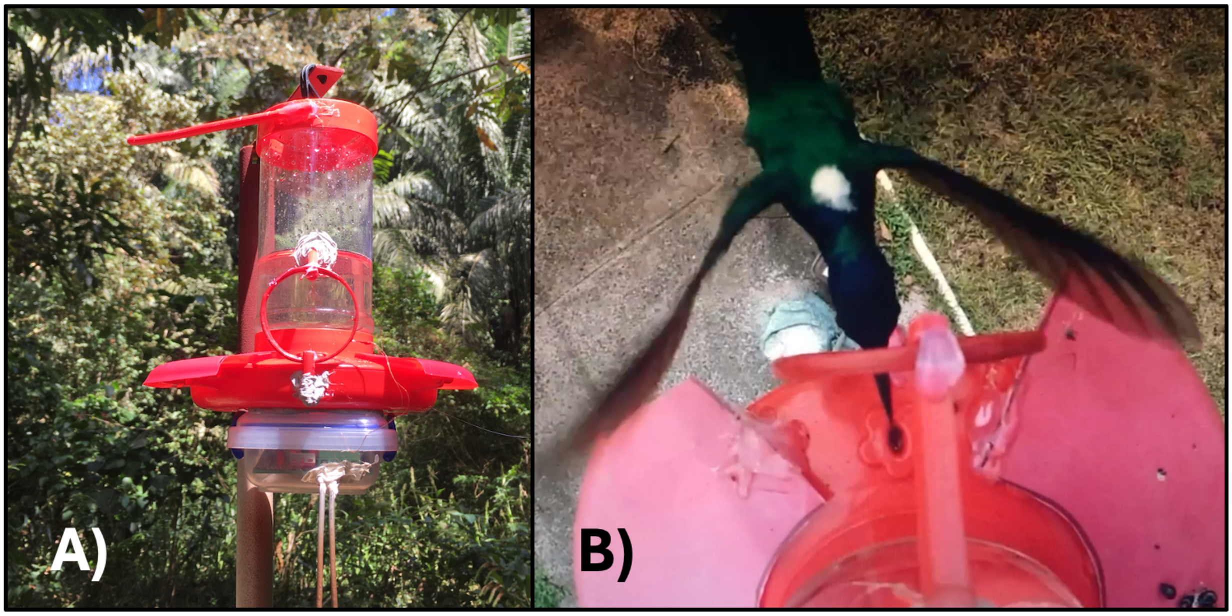


Supplemental Figure S2: Hummingbird feeders in this study were equipped with an RFID antenna which logged the presence of small tags implanted under the skin of white-necked jacobins during feeds. First Nature Hummingbird feeders (993051-001) were altered by removal of the perching ring surrounding the base so that birds could not perch next to the antenna. During prototype testing, antenna would detect perched birds that were not feeding. Therefore, we removed the perch so that all records of presence indicated active feeding. Two small pieces of the perch were then cut and used to hold the antenna in place in front of the feeding port and were attached to the feeder using plumbing sealant. Antennae were held onto these pieces with small amounts of hot glue, which could be removed during feeder cleanings. A third, larger piece of the perch was reattached to the top of the feeder away from the antenna detection region, allowing birds to perch on the feeders without recording. Antennas (Qkits 5.5 cm Sku AN0101) were dipped in red Plastidip sealant for weather-proofing. Other feeding ports were blocked using a red plastic covers. Logging devices were housed in a sealed plastic container with two packets of silicone for to prevent moisture accumulation. Antenna leads were attached to the logger by passing under the lid. Wires to the battery were attached through two small holes drilled into the container, which were re-sealed using plumbing sealant after wires were passed through. Panel A shows a front view of an active feeder, and Panel B shows the top view of a white-necked jacobin feeding from an active feeder. RFID tags were inserted between the shoulders of the birds, just under the loose skin at the base of the neck. Birds access the feeding port through the circular antenna, creating an ideal configuration for the antenna to read the RFID tag.

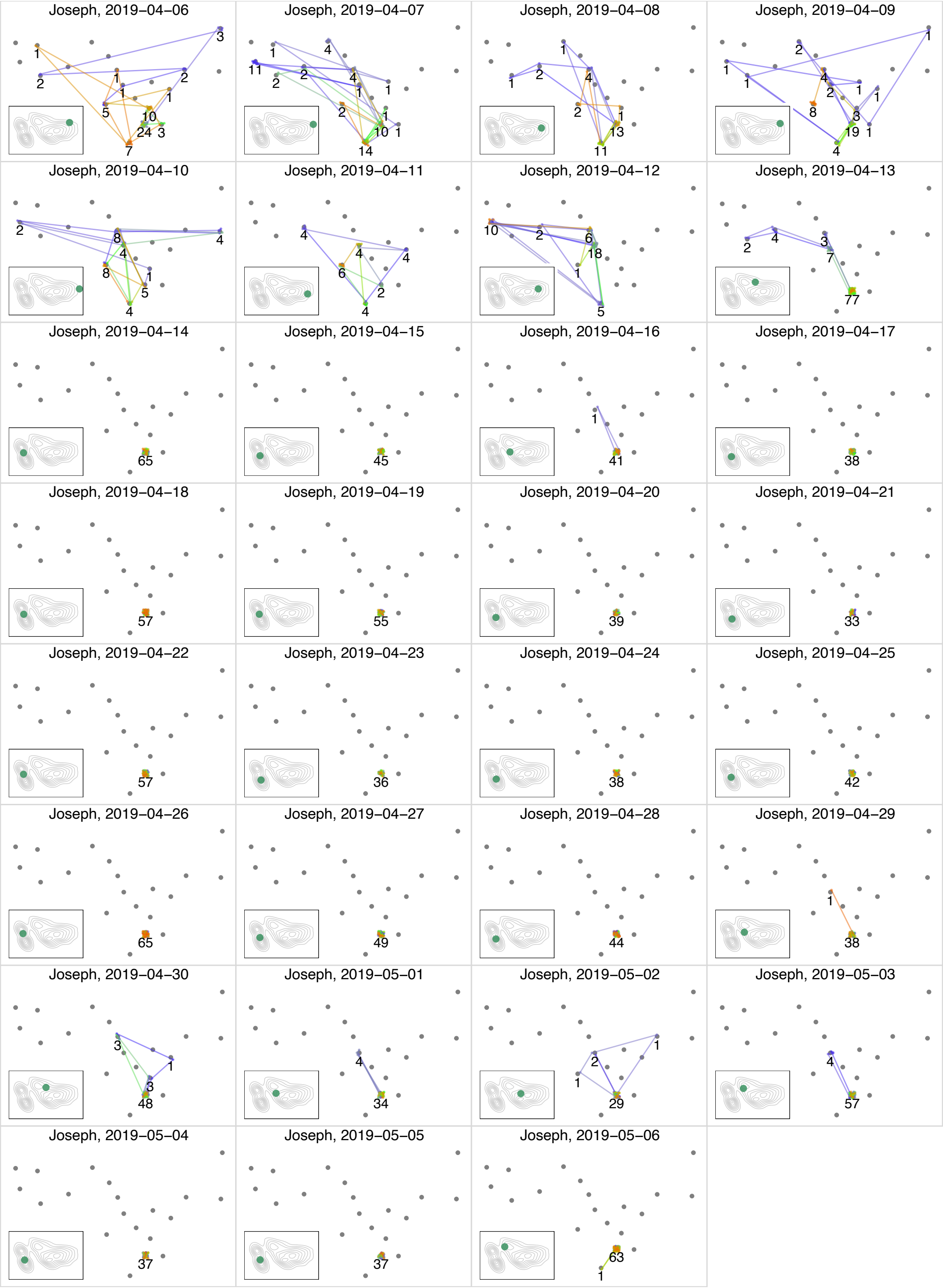


Supplemental Figure S2: One month of movement is shown for individual “Joseph” from April 6, 2024 to May 6, 2024. Grey points indicate available antenna-equipped RFID feeders. Points show feeds; lines show movement between feeders. Color spectrum indicates time of day, with purple representing mornings, green midday, and red late afternoon. Number of total daily visits to each feeder is indicated. Insets show the placement of each day within movement space (i.e., Figures 4 & 5, main text).

Supplemental Methods:

Birds were captured in Gamboa, Panama and the surrounding areas by placing mistnets around hummingbird feeders, or with drop-traps baited with hummingbird feeders. During capture, a 5-15 uL sample of blood was taken from the tarsal vein for genetic sexing. Blood was stored at room temperature in vials containing 0.25 mL of 2% SDS lysis buffer. At the Cornell Lab of Ornithology, we extracted DNA from blood samples using QIAGEN DNeasy Blood & Tissue Kits. 2550F/2718R primers were used to amplify a gene on the sex chromosomes (Fridolfsson and Ellegren 1999) which we were able to separate on a 2% agarose gel. Some samples could not be amplified with this primer pair, so we also used 1237L/1272H primers (Kahn et al. 1998) with fluorescently labeled 1237L. We then used an Applied Biosystems 3730xl sequencer at the Cornell Bioinformatics Facility to do a fragment analysis on the resulting amplified fragments. Individuals that we were not able to sex were removed from the dataset.

During feeding, hummingbirds will often consume nectar in a non-continuous manner by quickly alternating between active nectar consumption and short flights nearby, usually within a meter of the nectar source. Birds typically cycle through these behaviors rapidly before leaving or perching nearby. However, treating each non-consecutive feeding incident as a separate visit inflates the number of visits. To avoid this, we opted to set a threshold duration where gaps of a shorter period are considered a part of the same visit, while longer gaps designate separate visits. As with Falk et al. (2021) and Falk et al. (2022) which used the same dataset, here we used a threshold of 7 seconds. To identify this threshold gap length, all gaps between feeds from all individuals were first compiled. One-second gaps accounted for a large majority of gaps (41.5%), and the cumulative explanatory power of additional gap durations decreased with each additional second. Above 7 seconds, the increasing explanatory power of adding more seconds to the gap never exceeded one second. In Falk et al. (2021), results were not spurious to this threshold. While somewhat arbitrary, based on our observations a 7 second threshold balances the number of visits and the duration of each visit in a biologically relevant manner.
